# Phylogenomics reveals multiple introductions and early spread of SARS-CoV-2 into Peru

**DOI:** 10.1101/2020.09.14.296814

**Authors:** Eduardo Juscamayta-López, David Tarazona, Faviola Valdivia, Nancy Rojas, Dennis Carhuaricra, Lenin Maturrano, Ronnie Gavilán

## Abstract

Peru has become one of the countries with the highest mortality rate from the current severe acute respiratory syndrome coronavirus 2 (SARS-CoV-2) pandemic. To investigate early transmission event and genomic diversity of SARS-CoV-2 isolates circulating in Peru, we analyzed a total of 3472 SARS-CoV-2 genomes, from which 149 ones were from Peru. Phylogenomic analysis revealed multiple and independent introductions of the virus mainly from Europe and Asia. In addition, we found evidence for community-driven transmission of SARS-CoV-2 as suggested by clusters of related viruses found in patients living in different Peru regions.

## INTRODUCTION

Severe acute respiratory syndrome coronavirus 2 (SARS-CoV 2) is an emerging betacoronavirus responsible for the coronavirus disease 2019 (COVID-19) originated in December 2019 in Wuhan, China (1). Since then, the virus rapidly spread across the globe and was declared a pandemic on 11 March 2020 (2). Immediately, Peruvian government has carried out a series of sanitary interventions to prevent disease transmission including schools’ closure (11 March 2020), travel restrictions from Europe and Asia (12 March 2020), country borders closure (16 March 2020) and nationwide curfew (18 March 2020) (3). Despite these measures, SARS-CoV-2 has managed to rapidly spread causing over 702,776 cases and 30,236 deaths being the Lima city one of the major epicenters of SARS-CoV 2 infections in Peru, as of 11 September 2020 (4). As consequence, Peru has become one of the countries with the highest mortality rate from the current COVID-19 pandemic, although it is likely because of increase of diagnostic capacity and reported cases compared to other countries. Whole-genome sequencing (WGS) data of SARS-CoV-2 has rapidly become publicly available to have revealed insights about the genome structure as well as the temporal evolution and global transmission of the virus (5). However, the sources of epidemic transmission and genomic diversity of SARS-CoV-2 strains circulating in Peru to have led to the rapid spread of the virus remains poorly investigated.

We analyzed a total of 3472 SARS-CoV-2 genomes, from which 149 ones were from Peru to investigate how this novel virus became established in the country and to dissect the spread of the one in this area, which will help to orientate effective prevention measures to control the COVID-19 epidemic Peru.

## RESULTS AND DISCUSSION

Peru is one of the countries in the world with high levels of contagion and deaths by COVID-19 pandemic despite early national lockdown decreed by the government on 15 march when the country just had 71 cases reported. We sequenced 96 SARS-CoV-2 genomes obtained from patients with confirmed COVID-19 diagnostic up to July 04 at the National Institute of Health – Peru, yielding 87 high coverage-quality genomes. We removed 9 sequences with low coverage or too many private mutations/ambiguous sites indicative of sequencing error. These 87 cases were drawn from 7 Peru regions including Lima, Callao, Ancash, Lambayeque, Ica, Arequipa and Junin. Additionally, to these sequences, we have got 62 Peruvian SARS-CoV-2 genomes from GISAID database getting a genomic dataset of 149 Peruvian isolates from 12 regions that cover a period from the first official case on 05 march until 04-July (Figure 2A). Total Peruvian isolates were obtained from nasal and pharyngeal swab samples collected from 71 females (47.65%) and 78 (52.35%) males ranging in age between <25 years (14.37%), 25-35 years (17.36%), 36-46 years (22.65%), 47-59 years (21.56%), and >60 (24.06%) years.

Ninety-four (63%) out of 149 Peruvian sequences were obtained from Lima Metropolitana, the Peruvian capital, and 10 genomes from Callao, a neighbor city of Lima Metropolitana, where the Jorge Chavez International Airport is located and the only one in the country with flights to and from Asia, Europe, and North America (Figure 2A). Moreover, Lima-Callao is considered the epicenter of the Covid-19 pandemic for Peru presenting the 47% of total cases until 5 September (4). Our genomic dataset also contains 25 isolates from regions located in the Peruvian North (Lambayeque, La Libertad, Ancash, Cajamarca), 10 sequences from southern regions (Ica, Arequipa and Cuzco), 05 genomes from Loreto (the hardest-hit region by the pandemic in the Peruvian Amazon) and 05 from central regions of Peru (Junin and Huanuco) (Figure 2B).

We construct a time scaled phylogenetic tree from 3472 representative global SARS-CoV-2 genomes sampled between 24 December 2019 and 4 July 2020 including 149 Peruvian isolates (Figure 1A) to investigate early transmission event and genomic diversity of SARS-CoV-2 strains circulating in Peru. We found a strong temporal structure in our dataset by regressing genetic distance from tips to root in the ML tree against sample dates, resulting in a high correlation (r2 = 0.51) (Fig. S1). ML phylodynamic analysis revealed that the time to the most recent common ancestor (tMRCA) of the analyzed full genomic data was 28 November 2019 (20 November 2019 to 14 December 2020, CI 90%) being the inferred ancestral root Asia. These observations are in line with the known epidemiology of the pandemic (5). Furthermore, the analysis of genomic data has shown that Peruvian isolates were widely distributed across the phylogenetic tree suggesting multiple, independent introductions, designed as nodes (1–18) with over 70% of statistical support from ML (Figure 1A and Table 1). Similar to these introductions it has been reported in other countries such as Brazil (6), Colombia (7) and the USA (8). The most of these putative introductions of SARS-CoV-2 into Peru occurred between mid-February and early March, primarily sourced from Europe, Asia and South America (Table 1). Interestingly, the COVID-19 pandemic reached Latin America in February 2020 expanding into the region until March 2020 when the COVID-19 incidence curve started to grow more rapidly (3).

**Figure 1.**
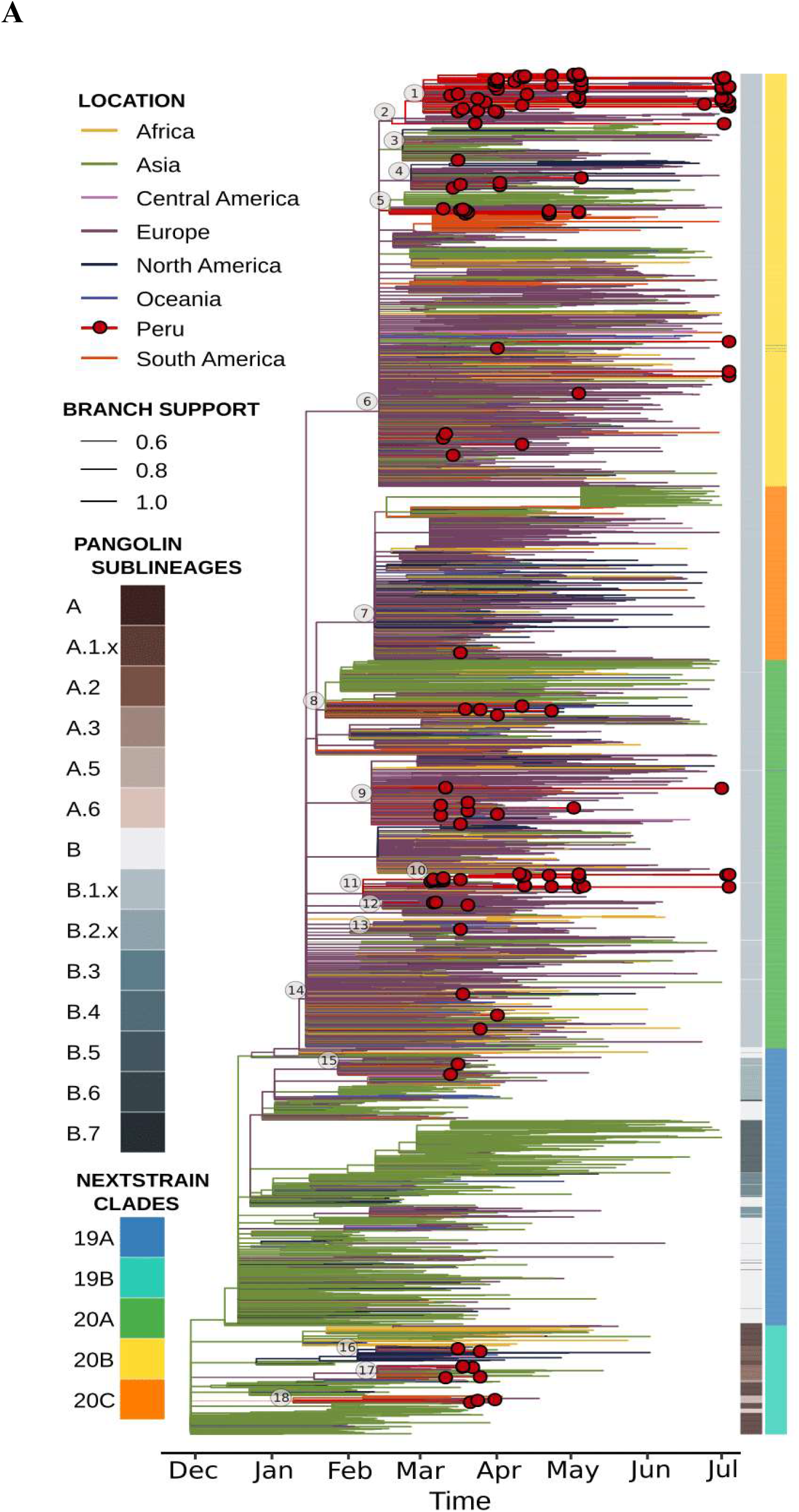
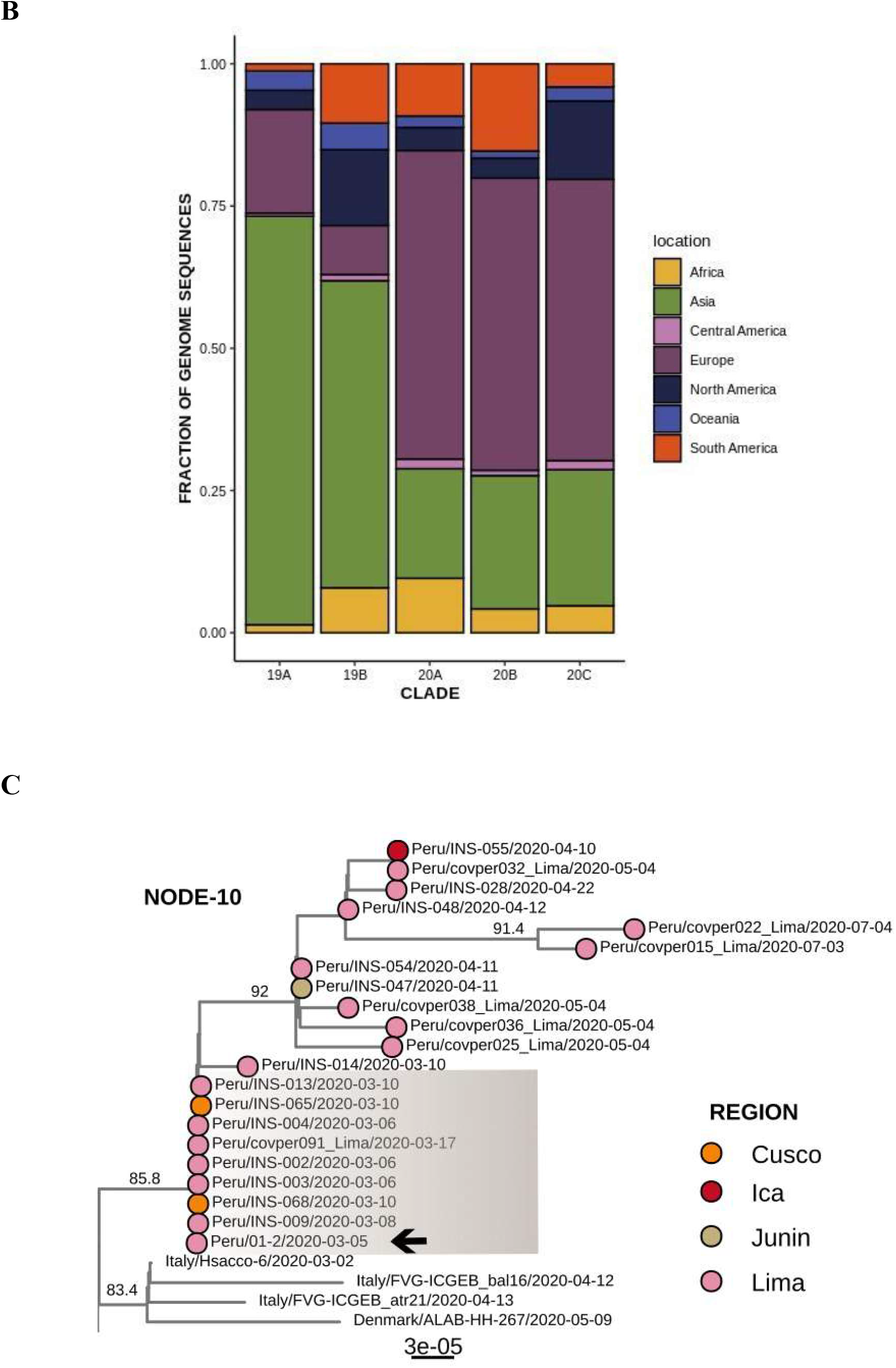
Phylogenetic relationships of SARS-CoV-2 from Peru and other global strains. **(A)** ML phylodynamic inference of 149 SARS-CoV-2 sequences from Peru in a global background of 3323 sequences available in the GISAID database as of 4 July 2020. Branches are colored according to the region of origin. Tip circles (red) indicate the position of the 149 Peruvian isolates. Clades that contain Peru sequences are highlighted, with names shown on the left. The node positions for the transmission events are marked by the numbers in white circles. **(B)** Stacked barplot showing the fraction of genome sequences per location by clade. **(C)** Local transmission clusters on the ML tree showing the source of cases by departments. Bootstrap support values ≥70% are shown; sibling clusters are collapsed for easier visualization. The mutations identified specific to the community transmission cluster are indicated (ORF1b: H604Y; S: Y119V). The scale bar at the bottom indicates the number of nucleotide substitutions per site.

Peruvian isolates (57%) were mainly clustered with clade 20B. This clade is largely composed of isolates obtained from patients with COVID-19 in Europe (51%), suggesting that introductions from Europe account for the majority of cases found in Peru between February and early March 2020 (Figure 1B and Table 1). Precisely, as of 12 March, 2020, the Ministry of Health reported imported cases in Lima from Spain, France and Italy (9).

We also identified within clade 20B, five putative introductions (node 1 to 5) where the most Peruvian isolates were interspersed without grouping by country or geographic region. Within this clade, we also identified SARS-CoV-2 Peruvian sequences that were reproduced in the time-scaled inference with mutations T1246I and G3278S in the ORF1a gene that distinguish clusters of sequences from Peru and elsewhere, suggesting a local transmission event that likely occurred in early March (tMRCA 95% CI: 27 February to 2 March) (Table 1, node 1, Figure 1A and Figure S1B).

Similar to isolates in the clade 20B, SARS-CoV-2 isolates positioned in the clade 20A were distributed throughout among isolates from multiple regions being mainly composed of European isolates (54%) (Figure 1 and Table 1) that is line to earliest sequences observed at the base of this clade (France, Russia and Czech Republic). Thus, it is highly likely to be sourced of European origin. Within this clade, we also identified a Peruvian cluster which includes the first official COVID-19 case in Peru (Peru/INS-01-02) identified in Lima City on 5 March 2020 and with travel history to France, Spain and Czech Republic (9). The SARS-Cov-2 sequence of this patient clustered with other Peruvian isolates (from 6-Mar to 04-July) conforming a “Peruvian cluster”, all of which were closely related with European isolates confirming its likely origin (Figure 1 and Figure S1B).

The tMRCA of this cluster (node 10) was estimated to be 4 March (25 February to 4 March, 90% CI) and contains two specific amino acid substitution: H604Y in the ORF1b gene and I119V in the spike protein (Table 1). What is more, within the cluster, we also observed 6 identical genomes to Peru/INS-01-02 reported in Lima until 17-March, suggesting local spread (Figure 1C and Figure S1B). Interestingly, two identical sequences to Peru/INS-01-02 reported in Cuzco, in southern Peru, on 10-March were identified, suggesting a regional introduction of SARS-CoV-2 derived from the first case from Lima (Figure 1C and Figure S1B). Although by March 18 had nationwide curfew, phylogenomic analysis revealed cases derived from the “Peruvian cluster” reported between April and July to Lima, Ica and Junin, suggesting an epidemic spread of SARS-CoV-2 into Peru likely by people local mobility (10). These results together with the nationwide widely distributed Peruvian isolates within clade supports the community spread likely originated from Lima.

For the remaining clades (19A, 19B and 20C), we inferred SARS-CoV-2 virus putative introductions to Peru between mid-December 2019 and early March 2020 (Figure 1A and Table 1). The clades 19A and 19B have mainly included Asian isolates (72% and 54%, respectively) suggesting that introductions from Asia account for the majority of cases found in Peru in the early pandemic period (Figure 1A, B and Table 1).

Our study also revealed some early introductions into Peru before the first case was reported (Table 1). That is likely to happen due to during this period the air travels were permitted while pandemic was happening. However, there is not epidemiological data which supported it.

All Peruvian sequences were classified into 9 PANGOLIN lineages: A.1, A.2, A.5, B.1, B.1.1, B.1.1.1, B.1.5, B.1.8 and B.2. The main lineages found in Peru were B.1. and B.1.1.1 grouping 97 (65%) of all 149 Peruvian sequences. Lima-Callao presents all nine lineages found in Peru, whereas the rest of regions harbor lineages B.1, B.1.1, B.1.1.1 and B.1.5 (Figure 2B). Additionally, until march we observed the 9 PANGOLIN lineages in Lima-Callao including A lineages but after April, only B.1.1.1, B.1.1, B.1 and B.5 ones were registered by WGS (Figure 1A). This spread of B sub lineages into Peru might be associated with the mutation D614G at spike protein, hypothesized that G614 providing the virus to be more infectious than D614 (11). These results are supported by the global epidemiological information that showed the predominance of the G614 variant after SARS-CoV-2 spread into Europe in late February and March. In addition, most introductions into South America countries (e.g. Colombia or Brazil) included European B lineages isolates (6,7,12). However, any other factors might be involved including uneven sampling, chance and epidemiology reasons (13).

**Figure 2.**
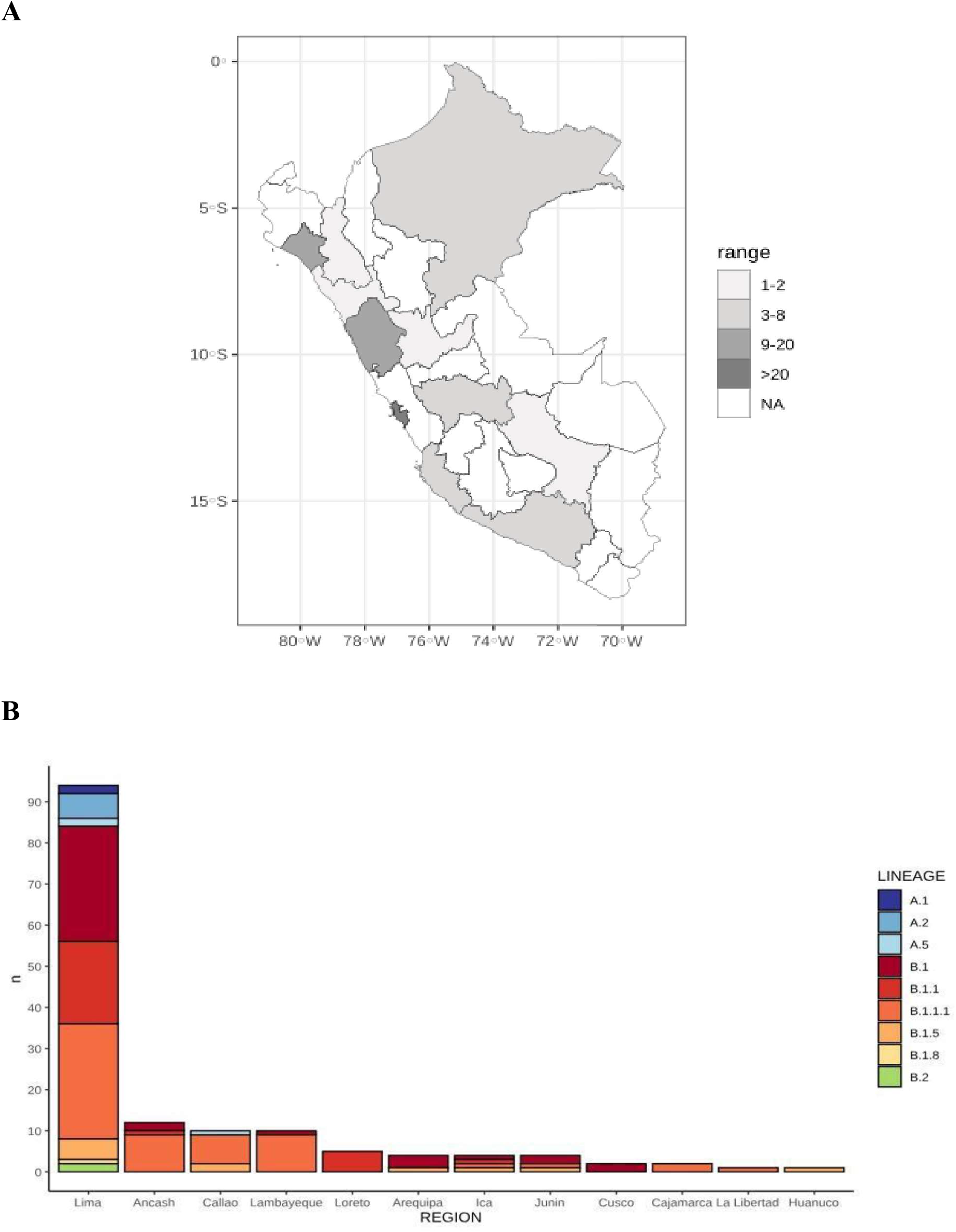
Geographic distribution of the Peruvian SARS-CoV-2 isolates circulating across regions of Peru. (A) Distribution of sequenced cases across Peru departments. (B) Breakdown of sequenced cases according to phylogenetic clades across Peru departments.

Furthermore, we identified non-synonymous mutations in all Peruvian SARS-CoV-2 isolates from sampled regions (Figure S2) of which the most frequent ones were in the D614G (S gene) and P314L (ORF1b). Interestingly, the D614G mutation has been related to increased infectivity (11) being also in linkage disequilibrium with the P314L mutation where both ones almost invariably co-occur (14). Therefore, that likely is one of the reasons for the viral rapid spread into all the country including Lima and Iquitos City (Loreto’s department) where the antibody prevalence has reached 25.3% and 67%, respectively, as of July 2020 (15).

One of the limitations of this study are the differences between the number of Peruvian isolates from cases identified within Peru regions. Specifically, there were more SARS-CoV-2 representatives’ sequences in Lima than in any Peru departments. Thus, it means that our estimates might be biased due to them being based on available background sequences at that time. Moreover, in the absence of epidemiological information such as travel history and contacts tracking, it is hard to associate periods of untracked transmissions with any specific regions or countries.

## CONCLUSIONS

We provided a first snapshot of the sources of epidemic transmission and genomic diversity of SARS-CoV-2 strains circulating in Peru in a global pandemic background, revealing multiple and independent introductions of the virus mainly from Europe and Asia. In addition, we found evidence that early spread of the virus in Lima City was sustained by community transmission. The data also underscore the limited efficacy of travel restrictions in a place once multiple introductions of the virus and community-driven transmission have already occurred. Our results emphasize the importance of using phylogenomic analyses to provide insights on sanitary interventions to limit the spread of SARS-CoV-2

Finally, we highlight the need for early and continued nationwide testing as well as the implementation of a real-time surveillance national system based on Whole-Genome Sequencing to identify untracked transmission clusters and to unscramble the evolution of the SARS-CoV-2 virus into Peru, and assess its impact on disease clinical severity.

## METHODS

### Sample collection and whole-genome sequencing of SARS-CoV-2

We obtained RNA of nasal and pharyngeal swabs from patients with confirmed COVID-19 diagnostic at the National Institute of Health - Peru (NIH-Peru), between 05 March 2020 and 04 July 2020 from departments of Lima, Callao, Ancash, Lambayeque, Ica, Arequipa, Junin. Samples that tested positive for SARS-CoV-2 RNA by RT-qPCR and with Ct < 25 were eligible to be included. The Genome-Whole Sequencing (GWS) of SARS-CoV-2 isolates (n=96) were performed on Illumina MiSeq at NIH-Peru using the CleanPlex® SARS-CoV-2 Panel (Paragon Genomics).

### SARS-CoV-2 genome assembly

The sequencing mean depth was 2,767.68. We generated an average reads number of 887,561 reads by sample. To remove low‐quality reads, trim off low‐quality we use fqCleaner v.0.5.0. Filtered reads were mapped against SARS‐CoV‐2 reference (NC_045512) using Burrows‐Wheeler Aligner MEM algorithm BWA‐MEM v0.7.7 (arXiv:1303.3997v2). SAMtools and Geneious Prime were used to sort BAM files, to generate alignment statistics and to obtain consensus sequence. Previous construction of the consensus genome sequence, we removed primer sequences with Software package fgbio using the primer genomic coordinates provided by Paragon Genomics.

### Peruvian and global collection of SARS-COV-2 genome sequences retrieved from GISAID

SARS-CoV-2 genomes from Peru (n=62) available in GISAID as of 26 August 2020 were selected until 07 April 2020 getting a dataset of 149 Peruvian genomes that cover a period from 05 March (officially first reported case) to 04 July of 12 Peruvian regions. In addition to this dataset, others countries sequences (n=3323) were retrieved randomly one genome per country per day (collection date) from GISAID between 26 December 2019 (including Wuhan-01 genome) and 01 July 2020 conforming a final genomic dataset of 3472 sequences to investigate the origins and genomic diversity of SARS-CoV-2 strains circulating in Peru by maximum likelihood phylodynamic approaches. The full genomic dataset was classified using the nomenclature of PANGOLIN (Phylogenetic Assignment of Named Global Outbreak LINeages) (16).

### Phylogenetic analysis of SARS-CoV in Peru

The full genomic dataset (n=3472) was aligned using MAFFT v 7.1 (10.1093/nar/gkf436) with default parameters. The alignment was manually curated, trimming the 5’ and 3’ ends and ambiguous regions. The final length of alignment was 29520 pb. We estimate a maximum likelihood tree of 3472 aligned sequences using IQ-tree v 1.6 (17) under a HKY nucleotide substitution model, with gamma distribution of among site rate variation (HKY+G+I) as selected by ModelFinder (18) and using the EPI_ISL_406801 sequence (GISAID) to root the tree. To measure branch support, we used Shimodaira-Hasegawa-approximate likelihood ratio test (SH-aLRT) with 1000 replicates. TempEst v1.5.1 (19) was used to assess the strength of temporal signal and inspect for outliers in the dataset by a root-to-tip regression of genetic distance against sampling date. We used TreeTime v0.7.6 (20) to estimate a time scaled phylogeny of ML tree. The analysis was performed using HKY substitution model and a coalescent Skyline prior to model population size changes under a strict clock (21). We also used the flag –confidence to retrieve node dates with 90% confidence intervals.

The clades were analyzed with NextStrain nomenclature for SARS-CoV-2 (https://nextstrain.org/ncov) Major clades that contained Peru sequences were defined by nucleotide substitutions and/or amino acid with ≥70% bootstrap support value in the ML tree and tMRCA with 90% of confidence intervals. To recognize local or introductions events were counted only for reproducible nodes with ≥70% bootstrap support value in the ML tree. Introduction events were defined as Peru sequences that clustered with non-Peru sequences across different clades. Local transmission events were defined as Peru-exclusive clusters with at least 5 or more sequences that were reproduced in the time-scaled inference (8). Statistical analyses and tree visualization using ggtree package were carried out in R version 3.6.2.

## ETHICS STATEMENT

This study was reviewed and approved by the Ethics Committee of the National Institute of Health from Peru (OI-011-20).

## AUTHORS AND CONTRIBUTORS

Conceptualization, E.J.L., D.T.; Formal analysis, E.J.L., D.C., D.T. and R.G., Methodology, E.J.L., D.C., D.T., F.V., N.R., and L.M.; Investigation, E.J.L., D.T., F.V., R.G.; Visualization, E.J.L., D.C., and D.T; Writing – Original Draft, E.J.L., and D.T.; Writing – Review & Editing, E.J.L., D.C., D.T., F.V., N.R., L.M. and R.G.; Supervision, E.J.L.

## ACKNOWLEDGMENTS

We are greatly grateful to all health personnel of Peru, especially to members of Respiratory Virus Laboratory from NHI-Peru for their dedication and courage in providing continued diagnostic and high-quality care under very difficult conditions.

The authors would like to thank Guillermo Trujillo and members of Respiratory Infectious Diseases Laboratory from NHI-Peru, especially to Helen Horna and Liza Linares for their support for the Whole-Genome sequencing.

The Research reported in this paper was supported by FONDECYT-CONCYTEC, under award contract number Proyecto N° 034-2020-FONDECYT.

The content is solely the responsibility of the authors and does not necessarily represent the official views of the NIH-Peru.

## REFERENCES

1. Wu F, Zhao S, Yu B, Chen Y-M, Wang W, Song Z-G, et al. A new coronavirus associated with human respiratory disease in China. Nature. marzo de 2020;579(7798):265–9.

2. Li X, Giorgi EE, Marichann MH, Foley B, Xiao C, Kong X-P, et al. Emergence of SARS-CoV-2 through Recombination and Strong Purifying Selection. bioRxiv [Internet]. 22 de marzo de 2020 [citado 10 de agosto de 2020]; Disponible en: https://www.ncbi.nlm.nih.gov/pmc/articles/PMC7255785/

3. Munayco CV, Tariq A, Rothenberg R, Soto-Cabezas GG, Reyes MF, Valle A, et al. Early transmission dynamics of COVID-19 in a southern hemisphere setting: Lima-Peru: February 29th–March 30th, 2020. Infect Dis Model. 1 de enero de 2020;5:338–45.

4. Ministry of Health. Sala situacional COVID-19 Peru [Internet]. 2020. Disponible en: https://covid19.minsa.gob.pe/sala_situacional.asp

5. Li J, Li Z, Cui X, Wu C. Bayesian phylodynamic inference on the temporal evolution and global transmission of SARS-CoV-2. J Infect. 2020;81(2):318–56.

6. Candido DS, Claro IM, Jesus JG de, Souza WM, Moreira FRR, Dellicour S, et al. Evolution and epidemic spread of SARS-CoV-2 in Brazil. Science. 4 de septiembre de 2020;369(6508):1255–60.

7. Laiton-Donato K, Arenas CJV, Ciro JAU, Munoz CF, Alvarez-Diaz DA, Villabona-Arenas LS, et al. Genomic epidemiology of SARS-CoV-2 in Colombia. medRxiv. 6 de septiembre de 2020;2020.06.26.20135715.

8. Gonzalez-Reiche AS, Hernandez MM, Sullivan MJ, Ciferri B, Alshammary H, Obla A, et al. Introductions and early spread of SARS-CoV-2 in the New York City area. Science. 17 de julio de 2020;369(6501):297–301.

9. Ministerio de Salud, Centro Nacional de Epidemiología, Prevención y Control de Enfermedades. Alerta epidemiológica ante la presencia de casos confirmados de COVID-19 en el Perú [Internet]. 2020 mar. Report No.: Alerta Epidemiológica código: AE-011-2020. Disponible en: https://www.dge.gob.pe/portal/docs/alertas/2020/AE011.pdf

10. Facebook. Facebook Data for Good to response to the COVID19 pandemic [Internet]. 2020. Disponible en: https://dataforgood.fb.com/docs/covid19/

11. Korber B, Fischer WM, Gnanakaran S, Yoon H, Theiler J, Abfalterer W, et al. Tracking Changes in SARS-CoV-2 Spike: Evidence that D614G Increases Infectivity of the COVID-19 Virus. Cell. 20 de agosto de 2020;182(4):812–827.e19.

12. Ramírez JD, Florez C, Muñoz M, Hernández C, Castillo A, Gomez S, et al. The arrival and spread of SARS-CoV-2 in Colombia. J Med Virol [Internet]. [citado 11 de septiembre de 2020];n/a(n/a). Disponible en: https://onlinelibrary.wiley.com/doi/abs/10.1002/jmv.26393

13. Grubaugh ND, Petrone ME, Holmes EC. We shouldn’t worry when a virus mutates during disease outbreaks. Nat Microbiol. abril de 2020;5(4):529–30.

14. Ogawa J, Zhu W, Tonnu N, Singer O, Hunter T, Ryan (Firth) AL, et al. The D614G mutation in the SARS-CoV2 Spike protein increases infectivity in an ACE2 receptor dependent manner. bioRxiv. 22 de julio de 2020;2020.07.21.214932.

15. Ministerio de Salud. Plataforma digital única del Estado Peruano [Internet]. 2020. Disponible en: https://www.gob.pe/minsa/

16. Rambaut A, Holmes EC, Hill V, O’Toole Á, McCrone JT, Ruis C, et al. A dynamic nomenclature proposal for SARS-CoV-2 to assist genomic epidemiology. bioRxiv. 19 de abril de 2020;2020.04.17.046086.

17. Nguyen L-T, Schmidt HA, von Haeseler A, Minh BQ. IQ-TREE: A Fast and Effective Stochastic Algorithm for Estimating Maximum-Likelihood Phylogenies. Mol Biol Evol. 1 de enero de 2015;32(1):268–74.

18. Kalyaanamoorthy S, Minh BQ, Wong TKF, von Haeseler A, Jermiin LS. ModelFinder: fast model selection for accurate phylogenetic estimates. Nat Methods. junio de 2017;14(6):587–9.

19. Rambaut A, Lam TT, Max Carvalho L, Pybus OG. Exploring the temporal structure of heterochronous sequences using TempEst (formerly Path-O-Gen). Virus Evol [Internet]. 1 de enero de 2016 [citado 11 de septiembre de 2020];2(1). Disponible en: https://academic.oup.com/ve/article/2/1/vew007/1753488

20. Sagulenko P, Puller V, Neher RA. TreeTime: Maximum-likelihood phylodynamic analysis. Virus Evol. enero de 2018;4(1):vex042.

21. Kingman JFC. The coalescent. Stoch Process Their Appl. 1 de septiembre de 1982;13(3):235–48.

