## Supplementary data for "Phylogenomics reveals multiple introductions and early spread of SARS-CoV-2 into Peru"

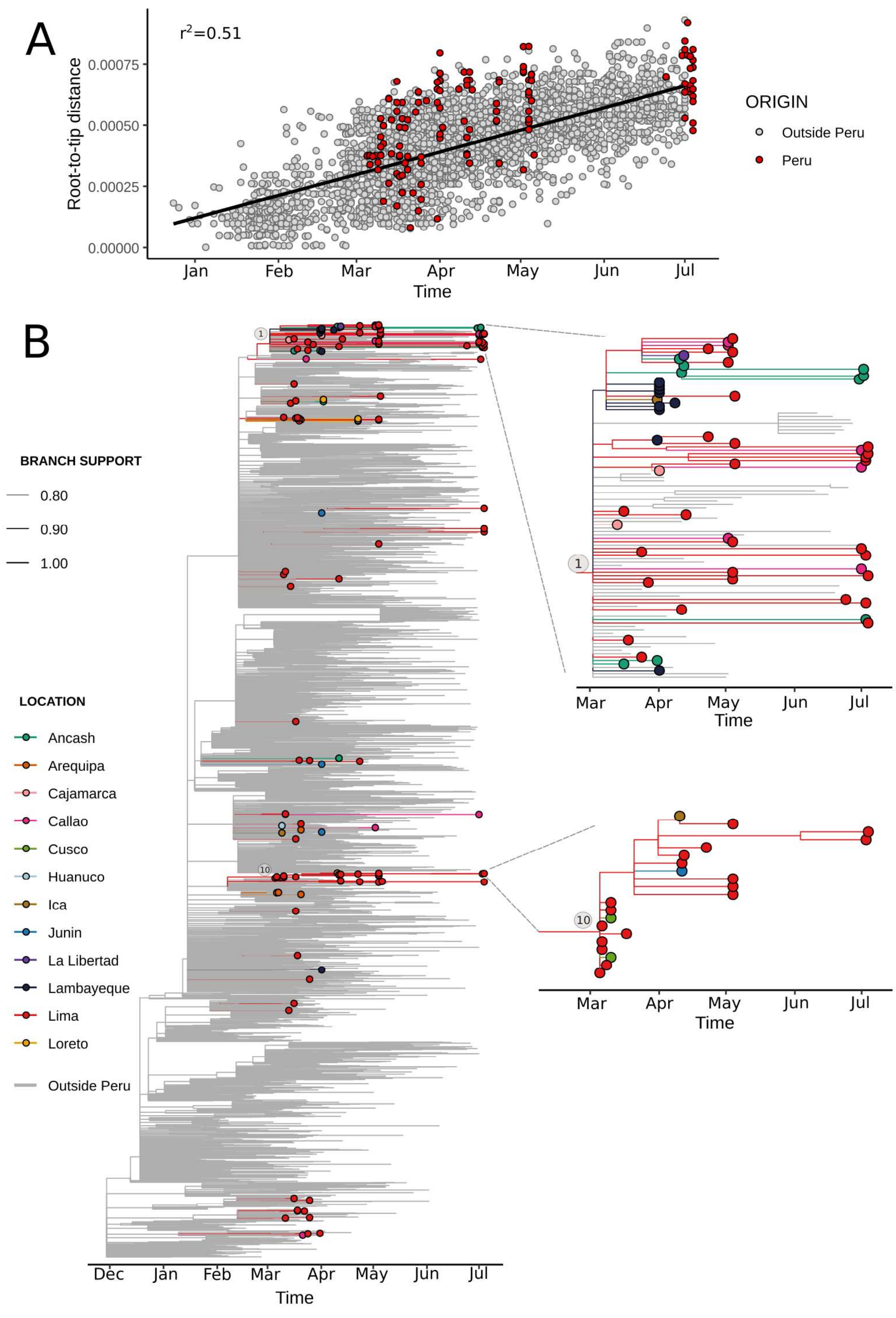

**Figure S1.** (A) Root-to-tip genetic distance regression of SARS-CoV-2 sequences against the dates of sample collection. Sequences are colored according to the geographic location: Peru and Outside Peru (B) ML tree scaled in time of 3742 SARS-CoV-2 genomes showing Peruvian sequences according to the region of collections. Branches colors represent most probable inferred locations and tip points are colored according Peruvian regions. Two main clades that grouped Peruvian sequences are amplified for easier visualization. The branch thicknesses are sized in proportion to the most probable inferred locations

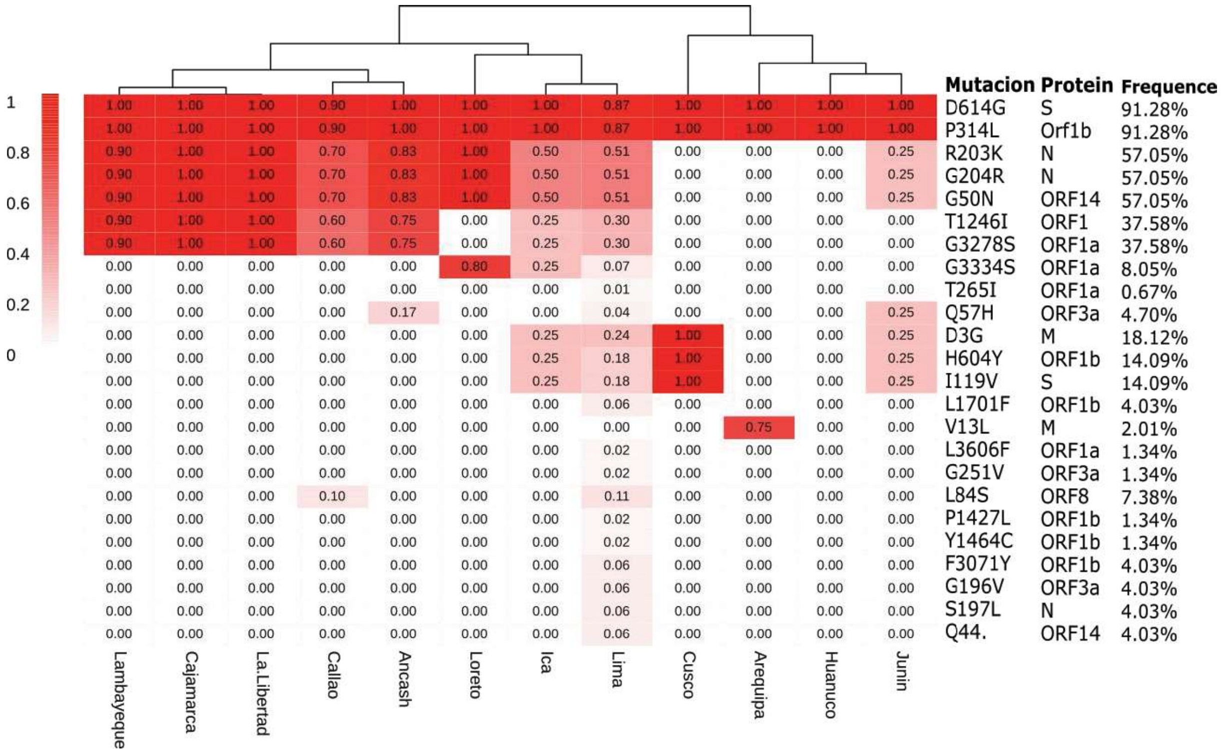


**Figure S2.** Heatmap of nonsynonymous amino acid substitutions of SARS-CoV-2 isolates circulating among 12 departments of Peru. The 12 departments were classified into three clusters based on the mutational signature by a hierarchical clustering. Protein sequence based on the SARS-CoV-2 (GenBank accession number NC_045512) is used as a reference.
